# Artifacts can be deceiving: The actual location of deep brain stimulation electrodes differs from the artifact seen on magnetic resonance images

**DOI:** 10.1101/2022.07.20.500820

**Authors:** Noa B. Nuzov, Bhumi Bhusal, Kaylee R. Henry, Fuchang Jiang, Jasmine Vu, Joshua M. Rosenow, Julie G. Pilitsis, Behzad Elahi, Laleh Golestanirad

## Abstract

**Introduction:** Deep brain stimulation (DBS) is a common treatment for a variety of neurological and psychiatric disorders. Recent studies have highlighted the role of neuroimaging in localizing the position of electrode contacts relative to target brain areas in order to optimize DBS programming. Among different imaging methods, postoperative magnetic resonance imaging (MRI) has been widely used for DBS electrode localization; however, the geometrical distortion induced by the lead limits its accuracy. In this work, we investigated to what degree the difference between the actual location of the lead’s tip and the location of the tip estimated from the MRI artifact varies depending on the MRI sequence parameters such as acquisition plane and phase encoding direction, as well as the lead’s extracranial configuration. Accordingly, an imaging technique to increase the accuracy of lead localization was devised and discussed.

**Methods:** We designed and constructed an anthropomorphic phantom with an implanted DBS system following 18 clinically relevant configurations. The phantom was scanned at a Siemens 1.5 Tesla Aera scanner using a T_1_MPRAGE sequence optimized for clinical use and a T_1_TSE sequence optimized for research purposes. We varied slice acquisition plane and phase encoding direction and calculated the distance between the caudal tip of the DBS lead MRI artifact and the actual tip of the lead, as estimated from MRI reference markers.

**Results:** Imaging parameters and lead configuration substantially altered the difference in the depth of the lead within its MRI artifact on the scale of several millimeters − with a difference as large as 4.99 millimeters. The actual tip of the DBS lead was found to be consistently more rostral than the tip estimated from the MR image artifact. The smallest difference between the tip of the DBS lead and the tip of the MRI artifact using the clinically relevant sequence (i.e., T_1_MPRAGE) was found with the sagittal acquisition plane and anterior-posterior phase encoding direction.

**Discussion/Conclusion:** The actual tip of an implanted DBS lead is located up to several millimeters rostral to the tip of the lead’s artifact on postoperative MR images. This distance depends on the MRI sequence parameters and the DBS system’s extracranial trajectory. MRI parameters may be altered to improve this localization.

## Introduction

Deep brain stimulation (DBS) is a neurosurgical procedure that involves implanting electrodes into specific brain regions and delivering electrical pulses via an implantable pulse generator (IPG) to improve symptoms of a variety of chronic neurological disorders. DBS may be used to improve quality of life for people with movement disorders such as Parkinson’s disease [1,2], essential tremor [3,4], and dystonia [5,6]. Furthermore, DBS is FDA-approved for treatment of epilepsy [7,8], severe obsessive-compulsive disorder [9-11], and is undergoing trials for treating major depression [12,13] and Alzheimer’s disease [14,15]. In some cases, DBS can be used at one of several targets to treat a single disorder. For example, improvements in specific Parkinson’s disease symptoms have been found when targeting either the subthalamic nucleus (STN) or the globus pallidus internus (GPi) [16,17].

Since the therapeutic effects of DBS heavily depend on the anatomical location of the implanted contacts within the target [18,19], it is crucial to assess the accuracy of different imaging modalities in determining the actual coordinates of an implanted DBS electrode. Valid electrode localization allows for the correct correlation of clinical outcomes with electrode position, as well as better compensation of any positioning error with changes in stimulation programming [20-23]. Magnetic resonance imaging (MRI) is commonly used for postoperative imaging of DBS electrodes thanks to studies addressing its safety [24-30]. However, because DBS leads contain metallic materials, significant susceptibility artifacts are apparent on MRI scans, hindering accurate electrode localization [31]. The artifact also has a radiofrequency (RF) component due to secondary magnetic fields induced on lead wires by MRI transmit fields, causing hypointense and hyperintense areas within the lead’s general artifact. The artifact is much larger than the lead itself [32,33], making it difficult to identify the precise locations of the contacts and the lead’s tip. Additionally, since the lead is implanted at an angle deviated from the MRI scanner’s main magnetic field, the artifact shape becomes more complex and has varying widths at different levels along the lead [34].

The most common method for DBS lead localization involves registering postoperative or intraoperative computed tomography (CT) images to preoperative MRI scans, using the caudal tip of the lead’s hyperintense artifact on CT images to represent the actual tip of the lead [35]. However, DBS localization using CT/MRI fusion compared to postoperative MRI alone, where the tip of the lead is similarly chosen as the caudal tip of the MRI hypointense artifact, has been inconsistent. Some studies have found the difference in the lead’s location between the two methods to be small enough to deem either method accurate, specifically ≤ 1 millimeters (mm) [36-39], while others have concluded that the difference between the two is too large to ignore, as high as 2.75 mm [34,40,41].

Additionally, the registration of CT to MR images introduces a registration error, which can vary significantly depending on the type of software used [39,42,43]. Most importantly, CT/MRI fusion does not adequately account for brain shift [44-46] and pneumocephalus [47], which usually occur after lead implantation. One study also identified that lead migrations could occur due to technical error during implantation and poor fixation of the DBS lead to the skull [48]. Due to these errors and discrepancies, a method for accurately determining lead position through postoperative MR images alone is highly desired.

The goal of this study was to quantify the distance between the actual location of the tip of a DBS lead and the tip of the MRI artifact under varying imaging conditions and device configurations. We performed experiments with an anthropomorphic phantom implanted with a commercially available DBS system to precisely determine the ground truth and assess the MRI artifact for different lead trajectories and pulse sequence parameters. To facilitate the translation of our findings, we used a clinically optimized T_1_MPRAGE sequence at 1.5 Tesla (T) to compare the effect of pulse sequence parameters on determining the actual location of the DBS lead tip. Finally, we devised a technique to better localize the tip of the DBS lead relative to the tip of the MRI artifact and quantified its accuracy.

## Materials and Methods

### Phantom Design and Construction

Experiments were performed in a head-and-torso human-shaped phantom representing an adult (length 65 cm, width 45 cm, volume 26 L). The design and construction of the anthropomorphic phantom is described elsewhere [49]. We also designed 3D-printed grids, posts, and an IPG holder to help position the DBS system in a well-controlled and stable configuration similar to clinical cases. To determine the true position of the lead, a custom structure was designed and fabricated, which held the lead at its center along with four capillary glass tubes at equal distances from the lead (shown in Fig. 1. a). The custom holder prevented bending of the lead post-implantation which is a common occurrence in patients. The glass tubes acted as MRI reference markers since their artifacts were minimally distorted, which was confirmed through statistical analysis. To assess the error due to variation of the markers’ artifacts, we calculated the Euclidean distance between each pair of opposite markers for every sequence and trajectory. The standard deviation of the distances was 0.2 mm, which was deemed small enough to not affect lead localization.

**Figure 1:**
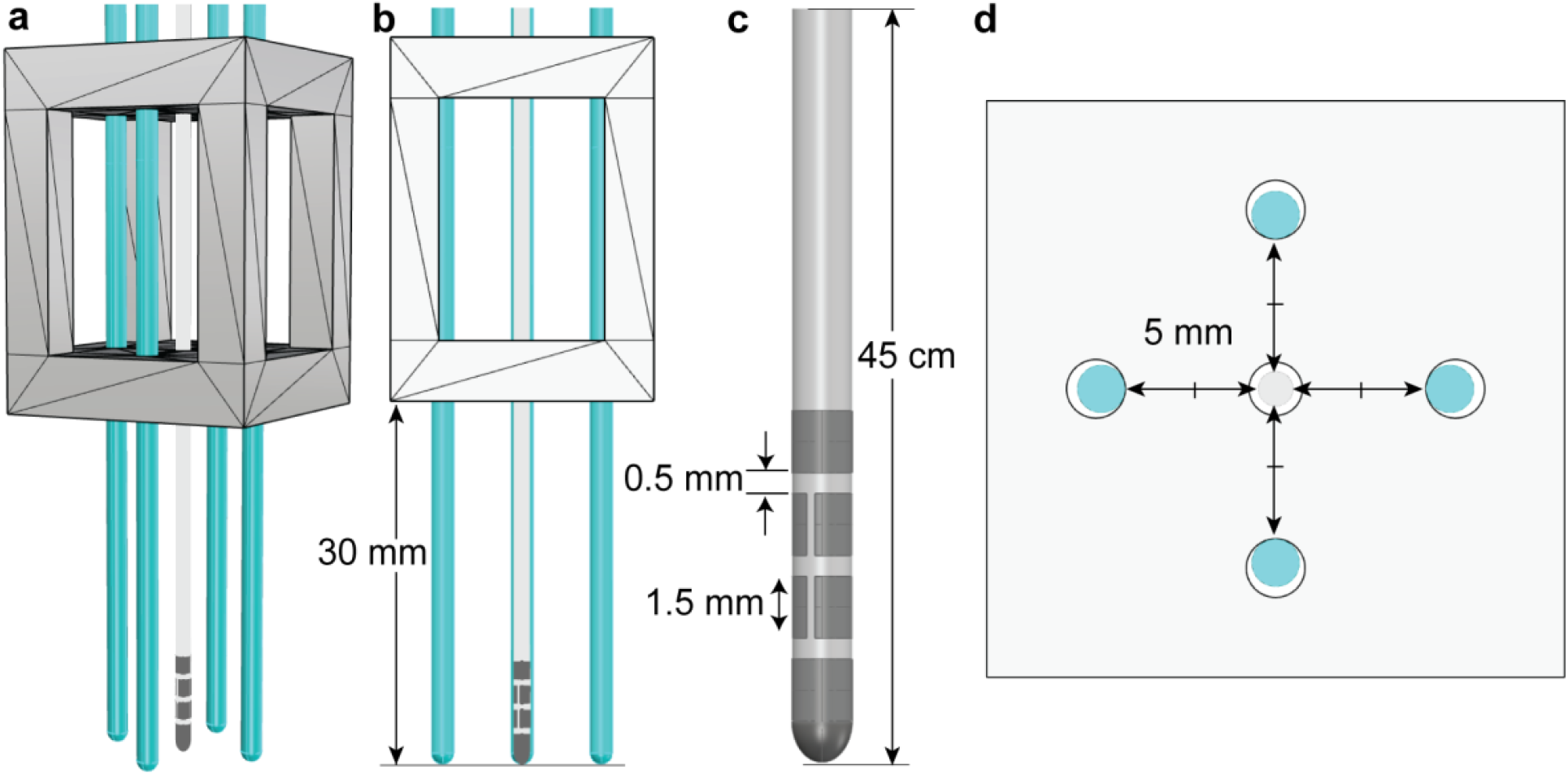
(a). 3D model of the DBS lead, custom holder (grey), and markers (blue). (b). Side view of the holder with lead and markers. (c). Boston Scientific DB-2202-45 directional lead used in experiments. (d). Top view of the holder with lead and markers.

Attention was paid to assure that the lead and markers were held completely straight and parallel, with the tips of all tube markers manually lined up to the same height as the tip of the lead, namely at 30 mm below the bottom of the holder (shown in Fig. 1. b). The holder held each marker exactly 5mm away from the lead in the plane perpendicular to the lead’s shaft (shown in Fig. 1. d). The lead and markers were attached to the inside of the phantom’s skull on the right side with a depth and penetration angle similar to what occurs in human patients (shown in Fig. 2. c). The electric conductivity of the medium surrounding the lead is shown to affect the field distortion and by proxy, the image artifact [26]. For this reason, we filled the skull with saline-doped agar solution with an electrical conductivity of σ = 0.34 siemens per meter (S/m) and relative permittivity of ε_r_ = 73.82, which is between the electrical properties of the brain gray and white matter. The rest of the phantom was filled with approximately 19 L of saline (2.6 grams NaCl per liter of double-distilled water) with electrical conductivity σ = 0.52 S/m and relative permittivity ε_r_ = 80.12, similar to the electrical properties of average tissue.

**Figure 2:**
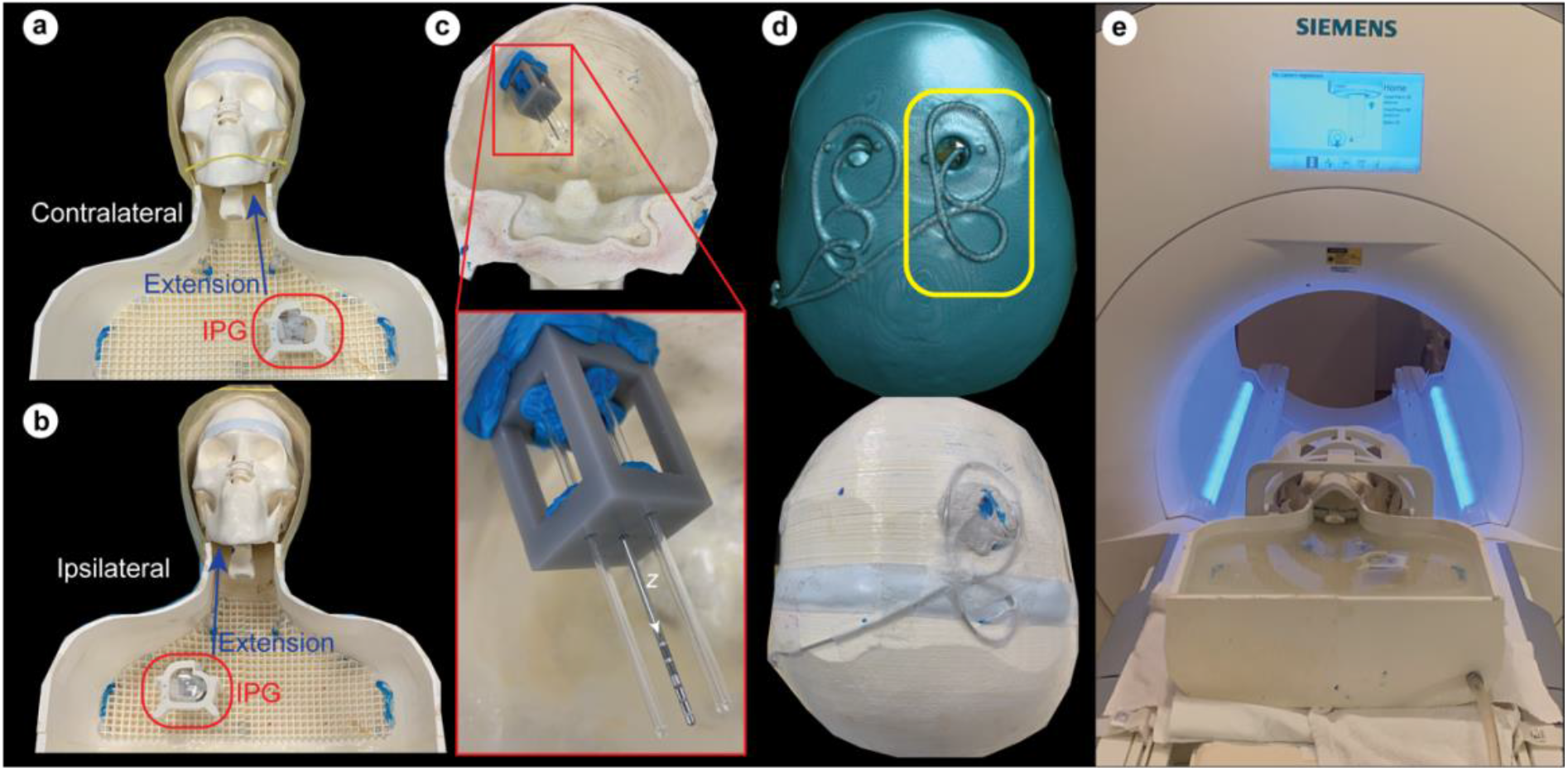
(a). Phantom setup for a contralateral extracranial trajectory. (b). Phantom setup for an ipsilateral extracranial trajectory. (c). Lead and marker holder attached to the inside of the right side of the skull. The z-axis is marked as the axis along the lead’s shaft. (d). Postoperative CT was used to segment trajectory 5 from patient (top) alongside the manually replicated trajectory used in the experiments (bottom). The right-sided trajectory was used (yellow box). (e). Phantom setup filled with saline entering the MRI scanner.

### DBS System Configurations

Experiments were performed with a 45 cm 8-contact directional lead (Boston Scientific, DB-2202-45, Vercise Cartesia™) (Marlborough, MA, USA) (shown in Fig. 1. c). The lead was connected to a 55 cm extension (Boston Scientific, NM-3138-55) which interfaced with the IPG (Boston Scientific, DB-1200, Vercise Gevia™). The IPG was positioned inside the holder which was then placed in the phantom’s pectoral region similar to the most common configuration in patients (shown in Fig. 2. a, b). A grid was placed inside the phantom to symmetrically alternate the IPG between the left and right side of the patient for contralateral (shown in Fig. 2. a) and ipsilateral (shown in Fig. 2. b) trajectories respectively. The extension was routed along the neck toward the IPG using posts.

The extracranial trajectory of DBS leads has been shown to affect their image artifact [49]. This is because lead trajectory affects the magnitude of RF-induced currents on internal wires [25,50-52], which causes the RF artifact (as well as RF heating). For this reason, we examined 18 clinically relevant lead trajectories (shown in Fig. 3) from which the first three were trajectories that exhibited moderate, minimum, and maximum RF heating at 1.5 T respectively, as determined by Bhusal et al. (2021) [49]. These included trajectories with (1) two concentric loops around the burr hole then routed towards the mastoid bone, (2) two concentric loops near the temporal bone then routed towards the mastoid bone, and (3) no loops on the skull and routed toward the mastoid bone (shown in Fig. 3). The latter 15 trajectories were derived from images of patients operated at Northwestern Memorial Hospital (n=7) and Albany Medical Center (n=8) (shown in Fig. 2. d). Trajectories 1 through 13 were contralateral and trajectories 14 through 18 were ipsilateral relative to the side of the IPG. For all trajectories, the excess of the lead extension was looped around the anterior surface of the IPG. The DBS system was programmed to MRI mode for the entire duration of the experiment.

**Figure 3:**
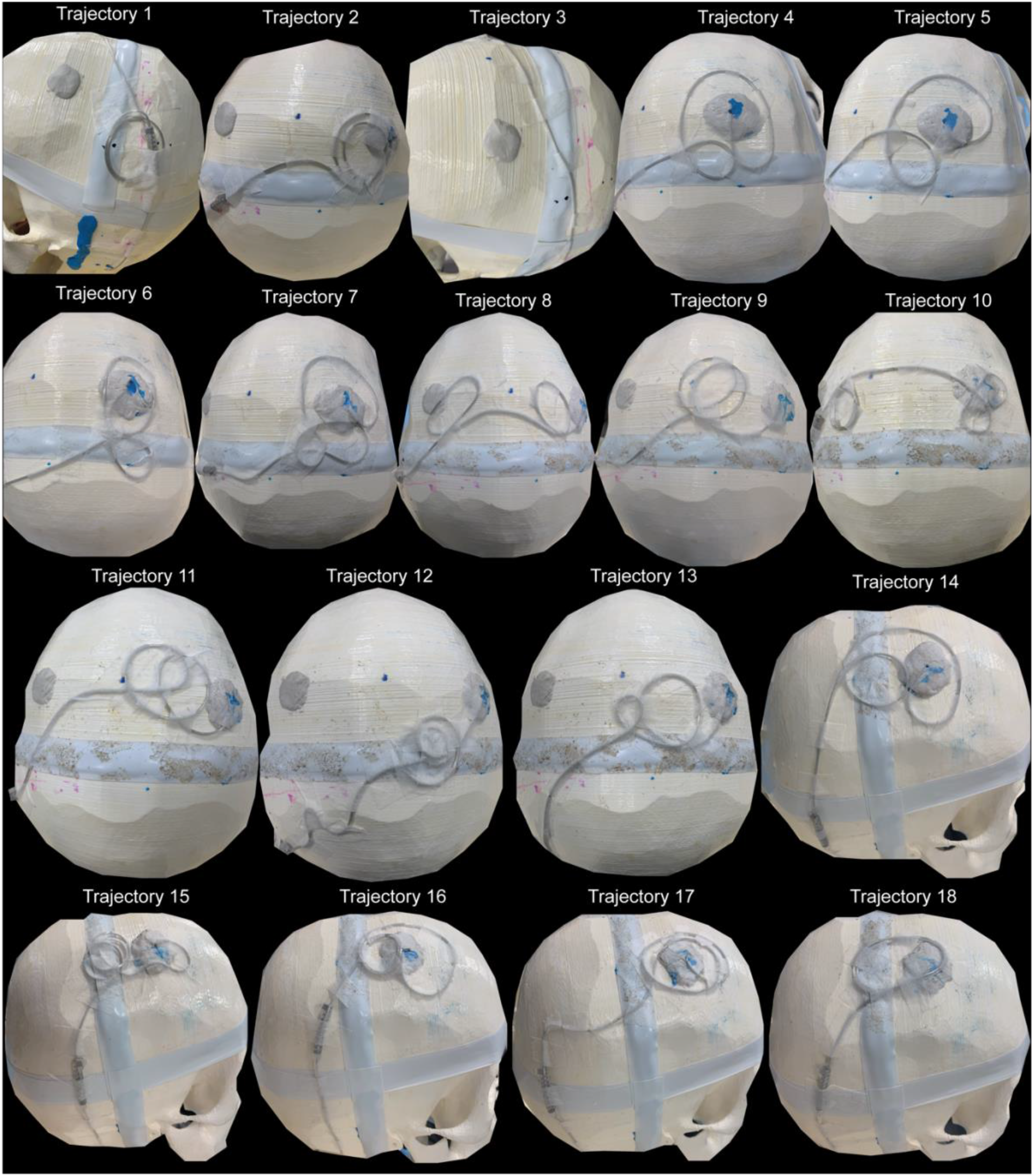
Photos of the extracranial trajectories 1-18 used in experiments

### MRI Sequences

The phantom was placed inside a 20-channel receive-only head/neck coil and scanned at a Siemens 1.5 T Aera scanner (shown in Fig. 2. e) using T_1_MPRAGE and T_1_TSE sequences with parameters given in Table 1. The T_1_MPRAGE sequence parameters were similar to those applied in the postoperative DBS workup of patients at Northwestern Memorial Hospital. In order to assess the effects of slice selection and phase encoding direction on the lead’s artifact, each sequence was repeated with axial (AX), sagittal (SAG), and coronal (COR) slice selection directions, as well as different corresponding phase encoding directions that did not cause aliasing (shown in Table 1). For the T_1_TSE sequences only, the axial scans were done perpendicular to the lead’s shaft while the sagittal and coronal scans were done parallel to the lead to illustrate unique patterning on the artifact.

**Table 1.**
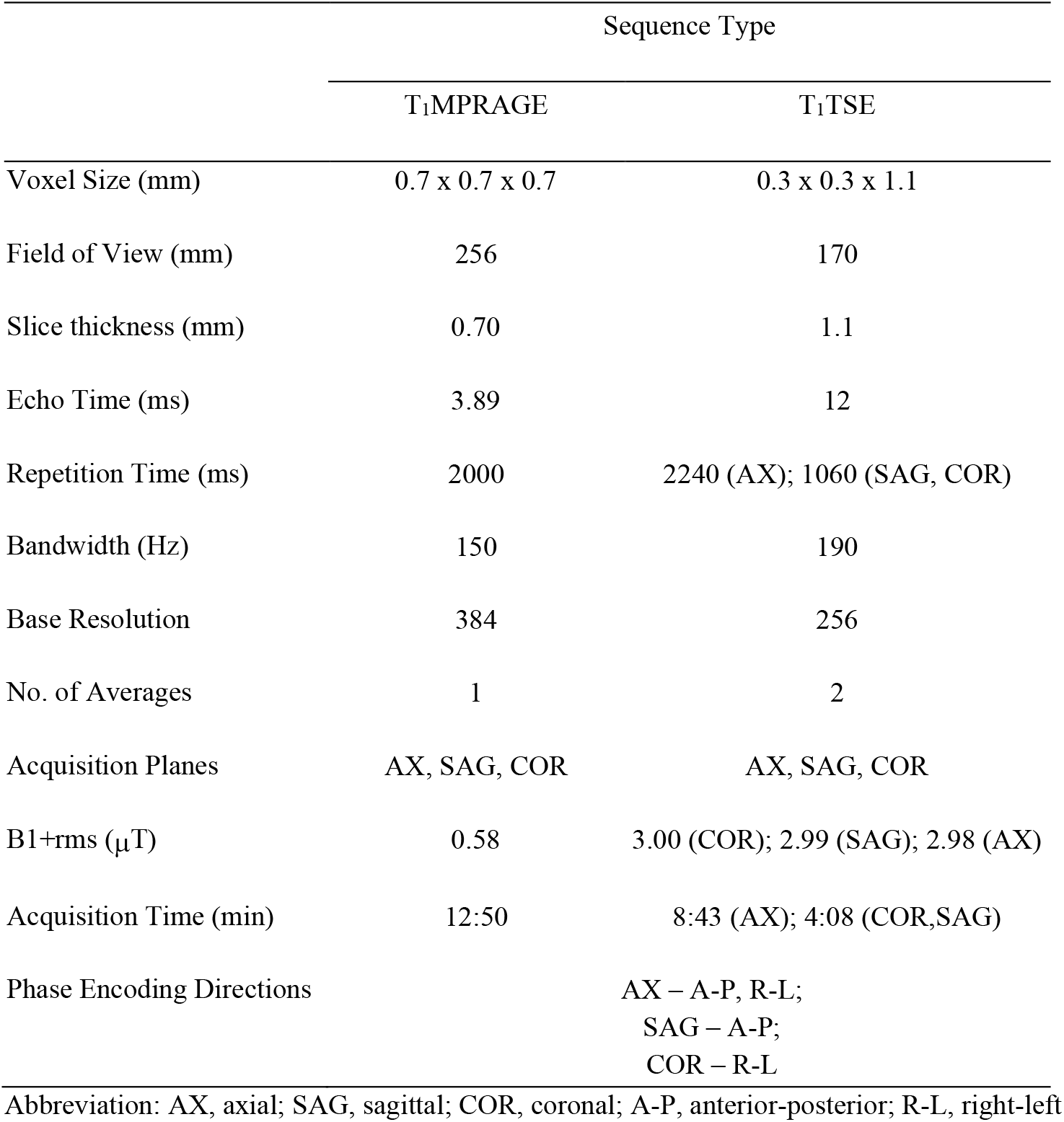
Parameters for MRI sequences at 1.5 T

### Image Segmentation and Artifact Analysis

MRI DICOM images were processed in 3DSlicer [53] where the *thresholding* tool was used to mask out the areas of hypointense signal (shown in Fig. 4. a). These masks were used to create 3D objects of the lead and marker artifacts which were then exported to a CAD tool, Rhino 3D [54], for further processing (shown in Fig. 4. b). Within Rhino 3D, we created individual center lines for the lead and marker artifacts and extracted the most caudal point of the centerline through the lead’s artifact to represent the “tip of the lead’s artifact” (shown in Fig. 4. c). To identify the ground truth, we first found the intersection of the two orthogonal planes that passed through the centers of each diagonal pair of markers and then selected the most caudal point of this line to represent the “actual tip of the lead” (shown in Fig. 4. d). For marker and lead artifacts that were highly distorted (e.g., trajectory 3) such that a centerline could not be made, the tips were chosen manually at the most caudal vertex of the artifact mesh. A line was connected between the actual tip of the lead and the tip of the lead’s artifact, *D* (shown in Fig. 4. e).

**Figure 4:**
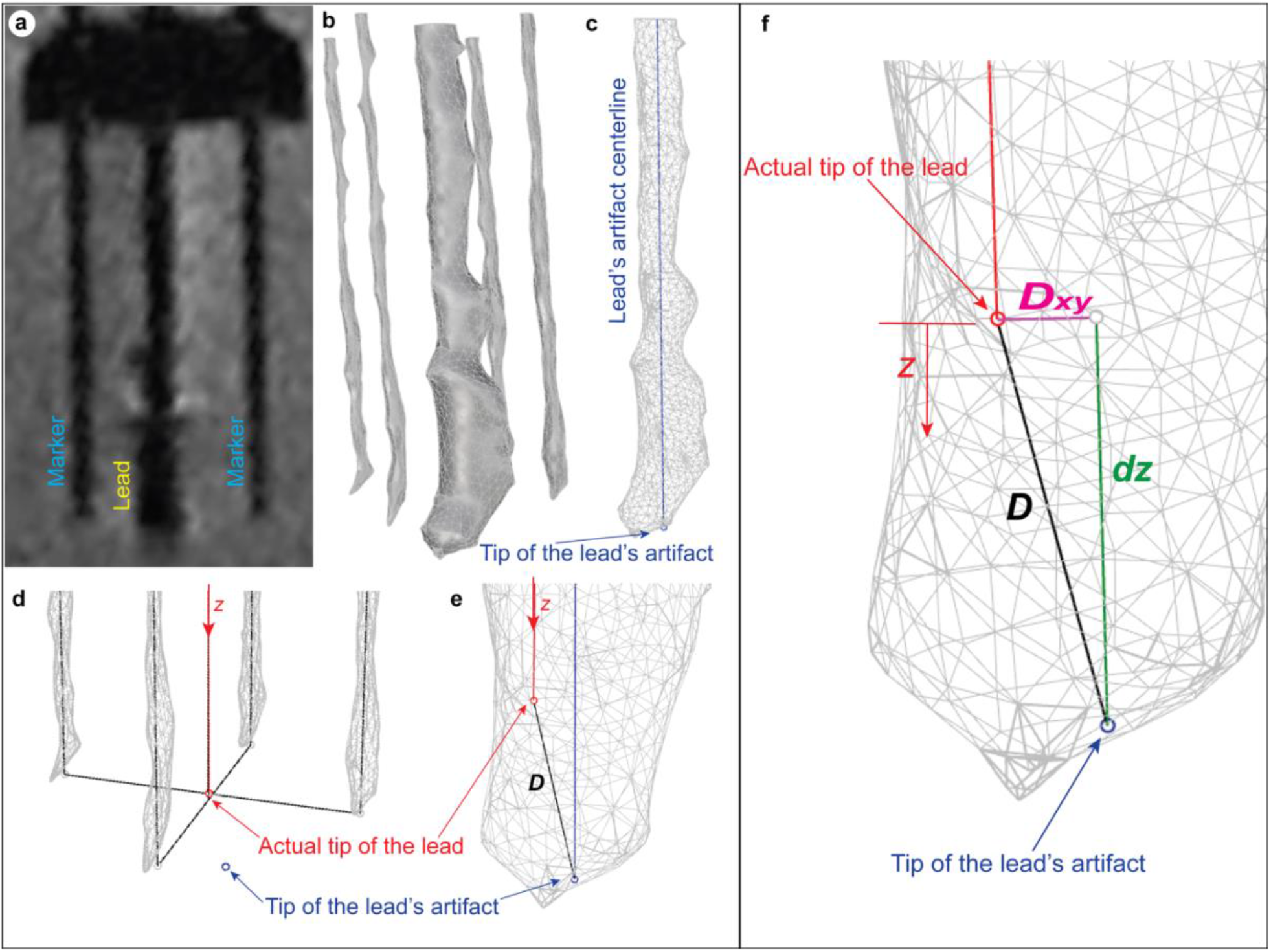
(a). MR image for trajectory 1, T1MPRAGE sequence, axial acquisition plane, anterior-posterior phase encoding direction. (b). Segmented 3D mask of the artifacts thresholded from 3D Slicer. (c). Mask of the lead’s artifact in Rhino (grey), centerline through the lead’s artifact (blue), and caudal point represents the tip of the lead’s MRI artifact (blue point). (d). Mask of the marker’s artifacts (grey), the intersection of the black planes represents the actual center of the DBS lead (red line) and the caudal tip of the DBS lead (red point). (e). Line D (black) between the actual tip of the lead (red point) and the tip of the lead’s MRI artifact (blue point). (f). Directional distance dz (green) from the actual lead tip to the tip of lead’s MRI artifact parallel to z-axis. Dxy (magenta) is line D between actual lead tip and lead’s artifact tip projected onto the plane perpendicular to lead’s shaft or z-axis.

To quantify the distance between the actual tip of the lead and tip of the artifact, we chose a coordinate system where the *z*-axis was aligned with the actual centerline of the lead and pointed through the lead’s shaft in the caudal direction (shown in Fig. 4. d, e). The Euclidean distance between the tip of the lead’s MRI artifact and the actual tip of the lead, *D*, was found (shown in Fig. 4. f). The directional difference in *z*-coordinates between the tip of the lead’s MRI artifact and the actual tip of the lead *dz* represented the depth of the MRI artifact in the caudal direction along the lead’s shaft below the tip of the lead (shown in Fig. 4. f). The perpendicular distance *D*_*xy*_ represents the magnitude of the projection of the line *D* onto the *x-y* plane perpendicular to the lead’s shaft (shown in Fig. 4. f). Microsoft Excel was used to calculate all distance formulas, averages, and standard deviations [55].

### Statistical Analysis

Distributions of the directional distance *dz* were grouped by sequence parameters, where there were 8 total sequences as given in Table 1 each repeated with n=18 trajectories. Due to normal underlying distributions but unequal variance, the Welch’s ANOVA was done to identify any significant differences in means between *dz* groups (*α* = 0.05). If any significance was detected for *dz*, the Games-Howell post-hoc test was applied to identify the significantly different pairs (*α* = 0.05). Statistical tests were not done with the Euclidean distance *D* and the perpendicular distance *D*_*xy*_ as these measurements are not directional and cannot be used to approximate lead location. All statistical tests were performed using the software R [56] run in RStudio [57].

## Results

Changing the sequence type, acquisition plane, phase encoding direction, and extracranial trajectory of the lead also substantially changed the directional, longitudinal distance *dz* between the actual tip of the lead and the tip of the lead’s artifact (shown in Fig. 5). In all cases, the tip of the lead’s artifact was more caudal along the axis of the lead’s shaft than the actual tip of the lead. MRI scans of all trajectories across all sequences are shown in Supplementary Figure 1.

**Figure 5:**
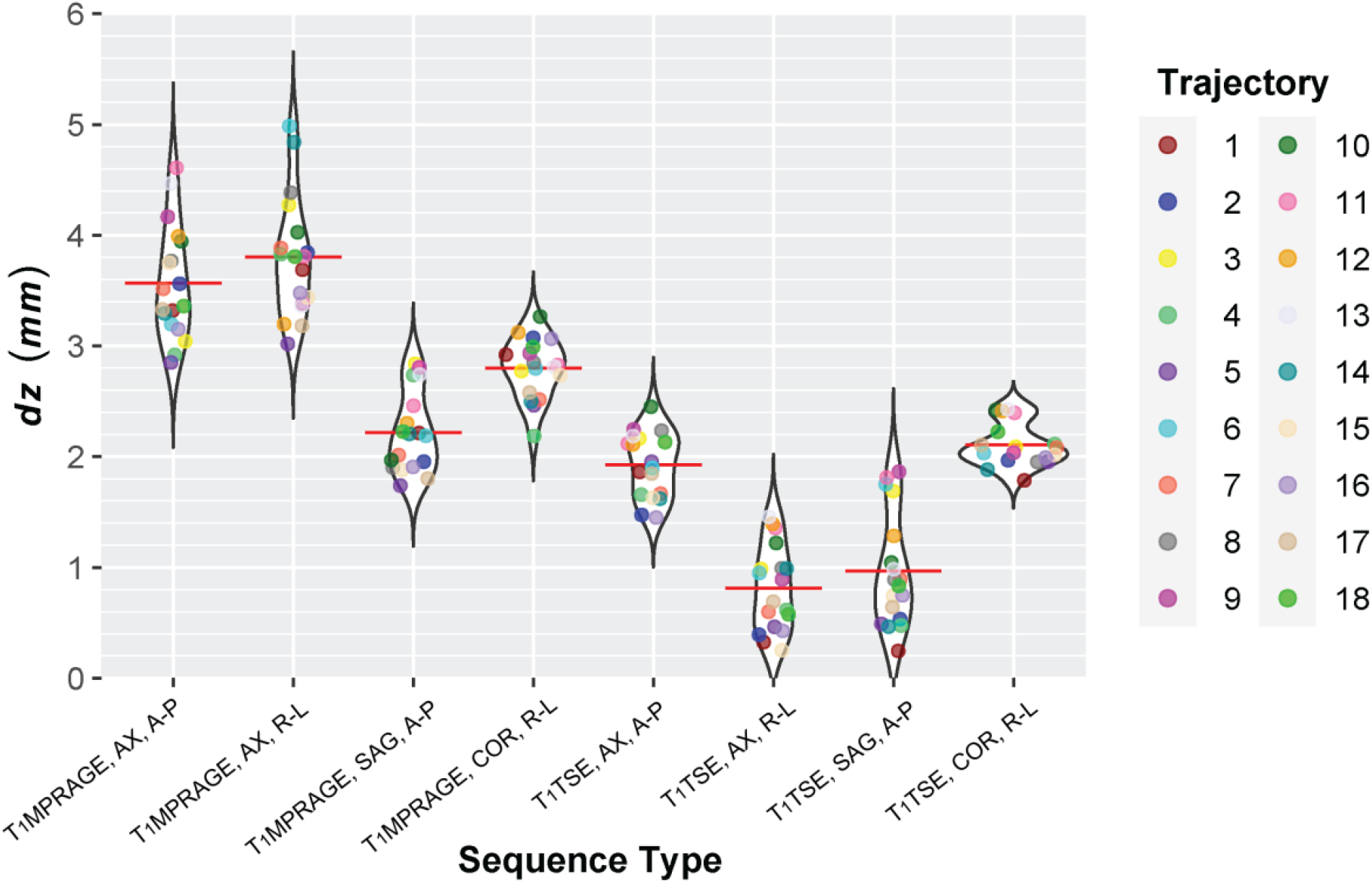
Directional distance dz for all trajectories and sequence parameters for T1MPRAGE and T1TSE in millimeters. Red lines represent the mean for each sequence. Trajectories 1-13 were contralateral and 14-18 ipsilateral. Trajectories 1-3 represent extreme SAR cases: (1) moderate heating, (2) least heating, (3) most heating.

Table 2 gives the mean ± standard deviation of *dz* (pooled over 18 trajectories). For the T_1_MPRAGE sequence, the largest average *dz* occurred for the axial acquisition plane and right-left phase encoding direction, while the smallest average *dz* was for the sagittal acquisition plane and anterior-posterior phase encoding direction. For the T_1_TSE sequence, the largest mean *dz* occurred for the coronal acquisition plane and right-left phase encoding direction, while the smallest mean *dz* for the axial acquisition plane and right-left phase encoding direction. For both T_1_MPRAGE and T_1_TSE sequences, the sensitivity to trajectory variation was minimized (i.e., the minimum standard deviation was observed in *dz*) when slice selection was in the coronal direction and phase encoding was in the right-left direction.

**Table 2.** Averaged directional distance 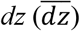 ± standard deviation in mm across all trajectories for each sequence.

|  | Sequence Type |  |
| --- | --- | --- |
|  | T<sub>1</sub>MPRAGE | T<sub>1</sub>TSE |
| AX, A-P | 3.569 ± 0.509 | 1.928 ± 0.295 |
| AX, R-L | 3.802 ± 0.547 | 0.810 ± 0.383 |
| SAG, A-P | 2.216 ± 0.362 | 0.966 ± 0.510 |
| COR, R-L | 2.801 ± 0.271 | 2.105 ± 0.195 |

Abbreviation: AX, axial; SAG, sagittal; COR, coronal; A-P, anterior-posterior; R-L, right-left

The mean value of *dz* for each respective sequence represents how far one must go rostrally along the lead’s shaft to identify the actual depth of the DBS lead tip (shown in Table 2). The difference between the individual *dz* values and the means for each respective sequence (i.e., *dz* – mean *dz*) was also found to demonstrate the range of error that would occur when using only the mean as the estimate of actual lead depth (shown in Fig. 6).

**Figure 6:**
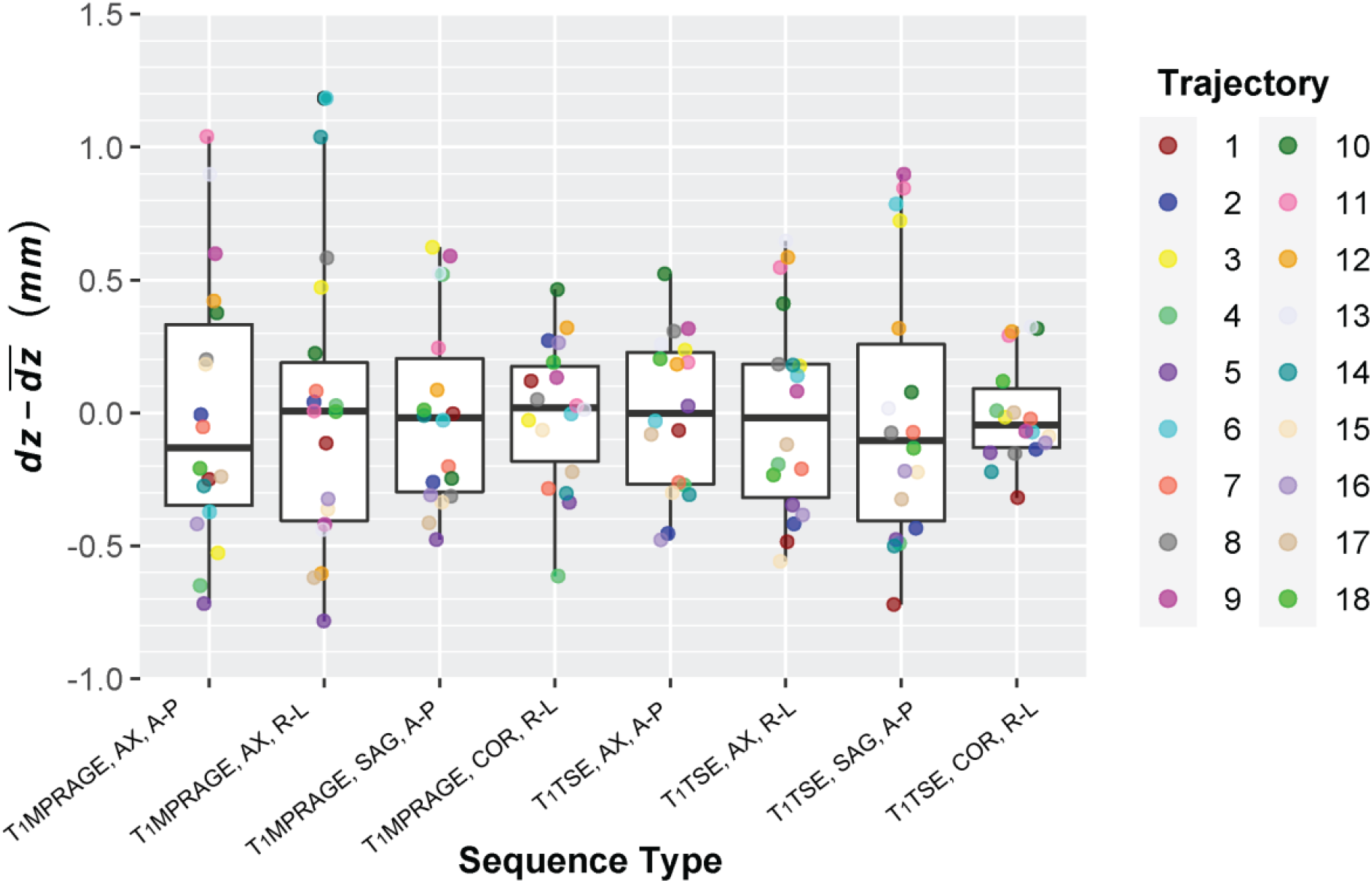
Difference between directional distances dz and the mean 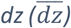 across all trajectories for each unique sequence in millimeters. Trajectories 1-13 were contralateral and 14-18 ipsilateral. Trajectories 1-3 represent extreme SAR cases: (1) moderate heating, (2) least heating, (3) most heating.

Welch’s ANOVA indicated that there was at least one pairwise comparison of mean *dz* with statistical significance (p << 0.001). Post-hoc analysis using G*Power calculated an effect size of 1.018 and power of 0.974 for the ANOVA [58]. The Games-Howell post-hoc test found that 23 out of 28 possible pairwise comparisons were statistically significant (p < 0.05) (shown in Table 3). Sequences that had the same acquisition plane and phase encoding directions, but different sequence type (T_1_MPRAGE vs T_1_TSE) had a difference in means that was statistically significant, which meant sequence type had an effect on the directional distance *dz*. For both T_1_MPRAGE and T_1_TSE, there were statistically significant differences between acquisition planes, even those with the same phase encoding direction. Lastly, for T_1_TSE, the axial acquisition plane scans were statistically different, demonstrating the significant effects of phase encoding direction (anterior-posterior vs. right-left).

**Table 3.** Results of Games-Howell post-hoc pairwise comparisons for mean 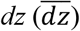 (***p<0.001; ****p<0.0001; ns, not significant).

| $\overline{dz}$ | | T<sub>1</sub>MPRAGE | | | | T<sub>1</sub>TSE | | | |
| --- | --- | --- | --- | --- | --- | --- | --- | --- | --- |
|  |  | AX, A-P | AX, R-L | SAG, A-P | COR, R-L | AX, A-P | AX, R-L | SAG, A-P | COR, R-L |
| T<sub>1</sub>MPRAGE | AX, A-P |  | ns | **** | *** | **** | **** | **** | **** |
|  | AX, R-L | ns |  | **** | **** | **** | **** | **** | **** |
|  | SAG, A-P | **** | **** |  | *** | ns | **** | **** | ns |
|  | COR, R-L | *** | **** | *** |  | **** | **** | **** | **** |
| T<sub>1</sub>TSE | AX, A-P | **** | **** | ns | **** |  | **** | **** | ns |
|  | AX, R-L | **** | **** | **** | **** | **** |  | ns | **** |
|  | SAG, A-P | **** | **** | **** | **** | **** | ns |  | **** |
|  | COR, R-L | **** | **** | ns | **** | ns | **** | **** |  |

Abbreviation: AX, axial; SAG, sagittal; COR, coronal; A-P, anterior-posterior; R-L, right-left

The values *D* represents the Euclidean distance, or length of the line, between the actual lead tip and the tip of the lead’s MRI artifact (shown in Fig. 7). All values of *D* were greater than zero, indicating a deviation of the artifact’s tip from the actual lead tip in 3D space.

**Figure 7:**
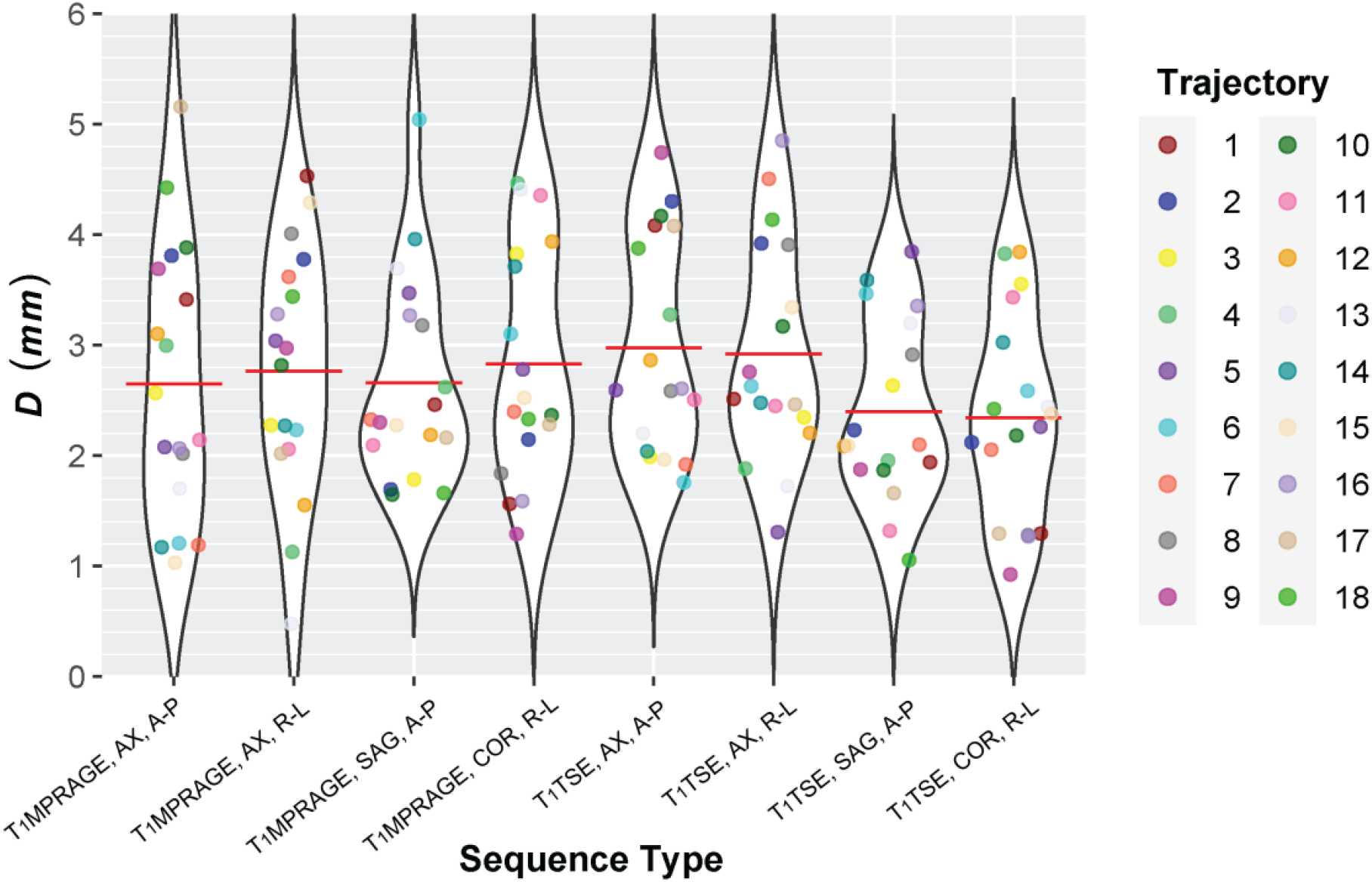
Length of the line D between the tip of the DBS lead and the tip of the lead’s MRI artifact across all trajectories and sequence parameters for T1MPRAGE and T1TSE in millimeters. Red lines represent the mean for each sequence.

The values *D*_*xy*_ representing the magnitude of the projection of line *D* between the actual lead tip and the artifact’s tip onto the *x-y* plane perpendicular to the lead’s shaft were also found (shown in Fig. 8).

**Figure 8:**
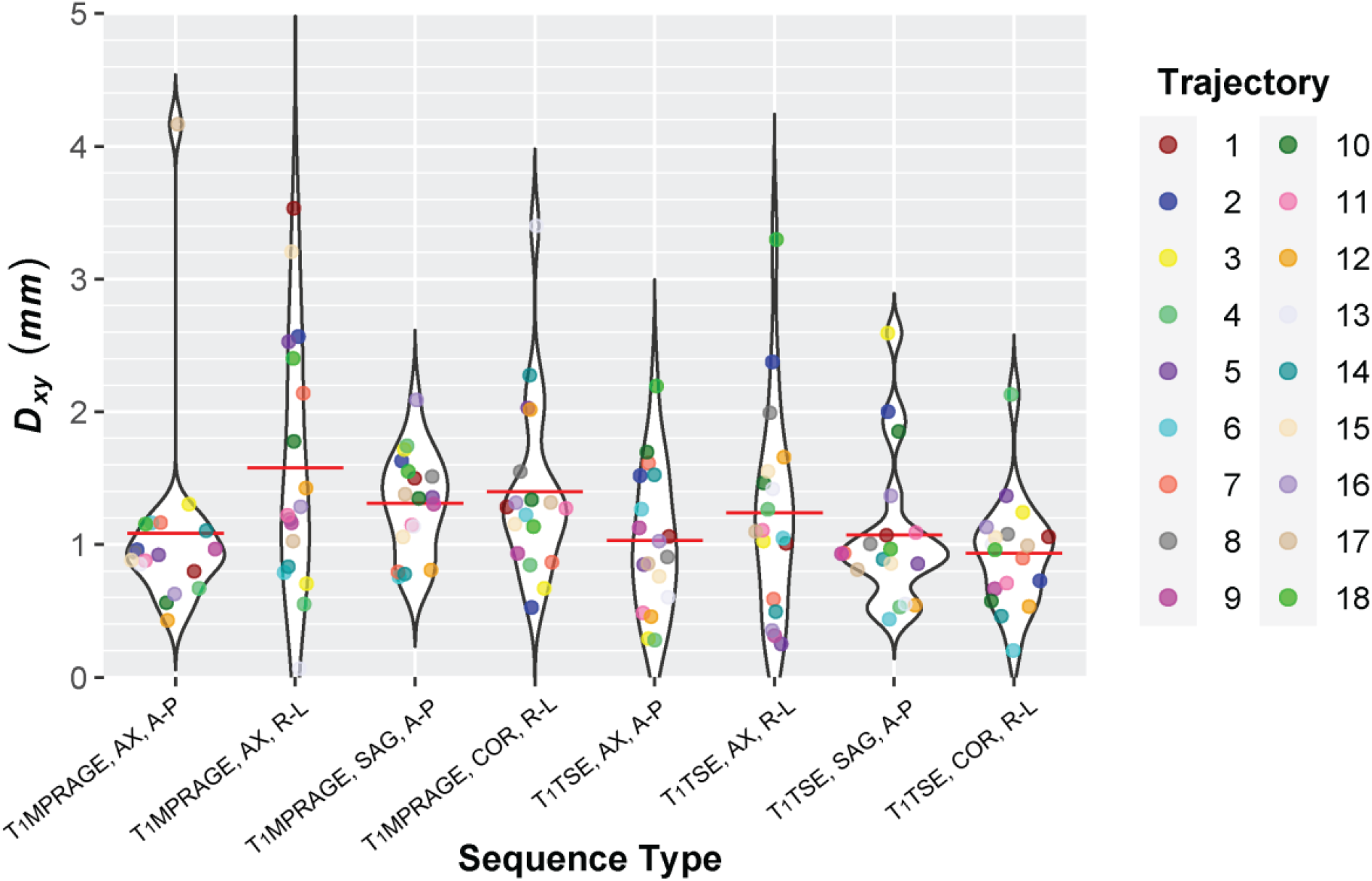
Magnitude of the perpendicular line D_xy_ across all trajectories and sequence parameters for T_1_MPRAGE and T_1_TSE in millimeters. Red lines represent the mean for each sequence.

The *D*_*xy*_ values were all greater than zero, indicating deviation of the artifact’s tip perpendicular to the *z*-axis or the actual lead’s shaft.

## Discussion/Conclusion

Current methods of determining the position of DBS electrodes from postoperative MR images rely on the assumption that the tip of the lead is located at the most caudal tip of the artifact [36,37,40]. However, researchers have recently begun formally disproving this assumption through phantom experiments [59]. To our knowledge, this is the first study identifying the difference in actual DBS lead positioning relative to its MRI artifact using a directional DBS lead system scanned at clinically relevant 1.5 T across multiple extracranial trajectories and varied MRI sequence parameters. We found that the actual tip of the lead is never located at the caudal tip of the MR image artifact, and the degree of variation along the lead’s shaft axis (i.e., rostral-caudal) changes with MRI sequence type, acquisition plane, phase encoding direction, and the extracranial trajectory of the DBS system.

One of the earliest papers investigating the MRI artifact of DBS leads found that the hypointense artifact extended 1.4 mm below the first contact of the Medtronic Activa^®^ 3389 lead (Medtronic, Minneapolis, MN) [33], while the lead itself has a 1.5 mm long plastic tip under the first contact which induces its own void artifact [42]. Furthermore, the same DBS lead was investigated in terms of its postoperative CT artifact, which only extended 1.2 mm below the bottom of the first contact [60]. This difference in the length of the postoperative CT and MRI artifacts demonstrates that selecting the tip of the artifact is not an appropriate method of localizing the lead across both modalities and also likely explains disparities in the CT/MRI fusion versus postoperative MRI localization research.

A study of patients who had their DBS leads removed suggested that the center of the hypointense MRI artifact can be assumed to justly represent the lead’s centerline in DBS patients [61]. This would allow for the determination of the mediolateral and anteroposterior coordinates of the lead, but the exact location of the lead’s tip in the rostral-caudal direction, *z*, was not determined [62]. A more recent phantom study investigated the distance between the tip of the actual lead and the tip of the lead’s MRI artifact and found differences using T1-weighted three-dimensional turbo-field echo (T1w-3D-TE) and T2-weighted turbo spin echo (T2w-TSE) sequences at 3 T with varying implantation angles; however, scans at this strength are not clinically used [59]. This study also disproved the previous determination [61] that the centerline of the lead’s shaft is aligned with the center of the lead’s MRI artifact. Furthermore, our non-zero values for the magnitude of deviation of the artifact’s tip in the plane perpendicular to the lead’s shaft, *D*_*xy*_, supports this newer conclusion.

Previous postoperative MRI studies have only been done using lead models with 4 ring-shaped omnidirectional contacts [33,34,40,41,59], while the investigation using directional or segmented electrodes is absent. So, our study employed an anthropomorphic phantom implanted with a directional DBS lead system where the effects of the extracranial trajectory, both contralateral and ipsilateral to the IPG, were investigated for the first time on the MRI lead’s artifact. With the ground truth known, we determined the distance *dz* parallel to the lead’s shaft between the actual tip of the DBS lead and the tip of the lead’s MRI artifact. The actual tip of the lead is always located in an anatomical position that is less deep or more rostral to the tip of the artifact. At a minimum, this distance *dz* along the lead’s shaft axis was 0.25 mm but varied up to 4.99 mm under certain conditions.

In this experiment, the T_1_MPRAGE sequence was based on routine scanning protocols of our DBS patients at Northwestern Memorial Hospital, where the axial acquisition plane and right-left phase encoding direction are used. The T_1_TSE sequence is valuable for contact localization since it demonstrates a unique alternating bright and dark intensity pattern representing the contacts and plastic insulation near the proximal end of the DBS lead (shown in Fig. 9. c). The unique patterning has been demonstrated on other types of sequences, such as T_2_TSE, and new sequences have also been recently explored that are optimal for DBS lead localization [63].

**Figure 9:**
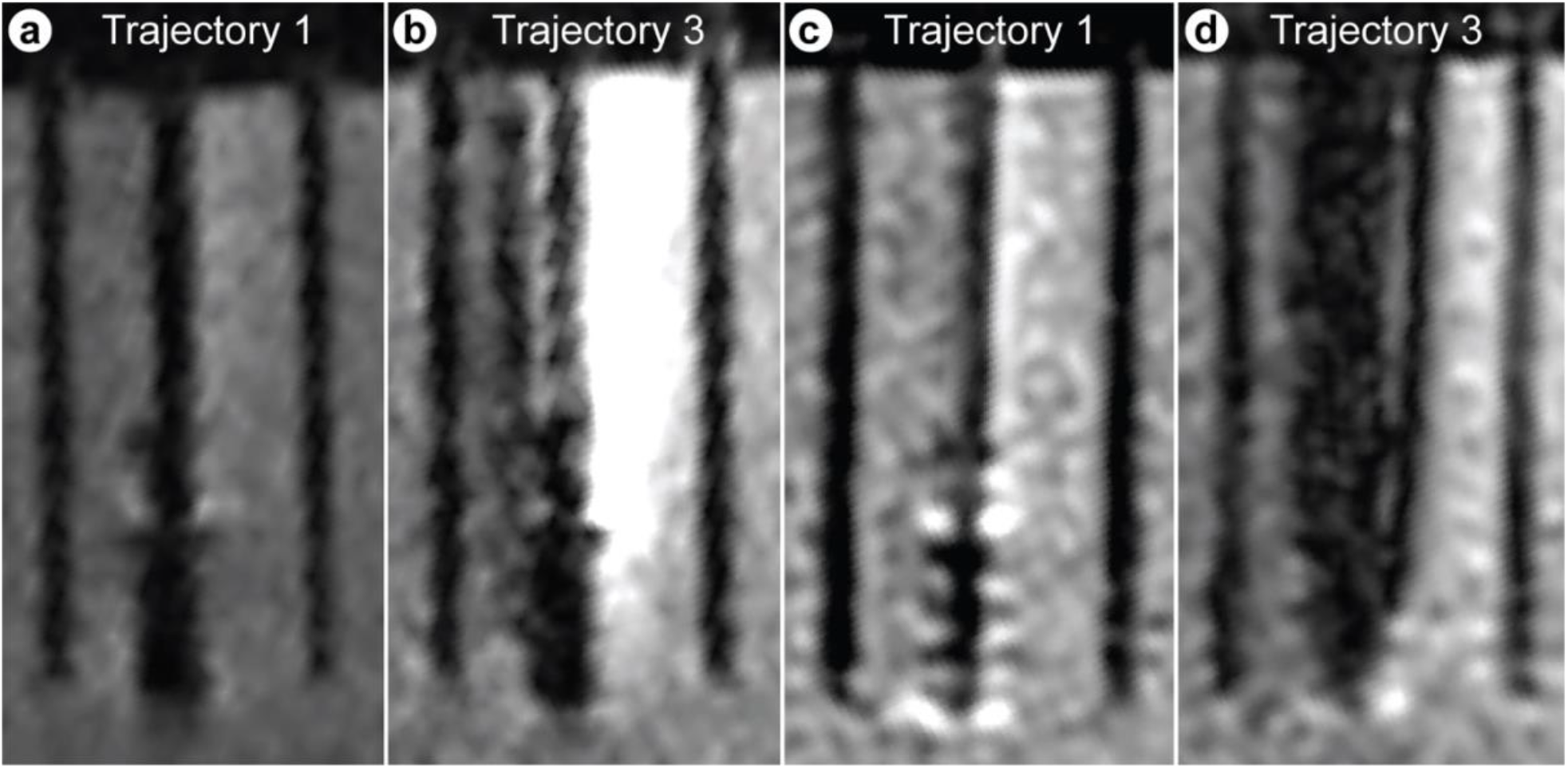
(a). T1MRAGE, axial acquisition plane, anterior-posterior phase encoding direction, scan of trajectory 1. (b). T1MRAGE, axial acquisition plane, anterior-posterior phase encoding direction, scan of trajectory 3. (c). T1TSE, sagittal acquisition plane, anterior-posterior phase encoding direction, scan of trajectory 1. (d). T1TSE, sagittal acquisition plane, anterior-posterior phase encoding direction, scan of trajectory 3.

We found that mean *dz* was the smallest using the sagittal acquisition plane and anterior-posterior phase encoding direction for T_1_MPRAGE, compared to the axial acquisition plane and right-left phase encoding direction for T_1_TSE. We also found that the artifacts and distances were affected by the extracranial trajectory of the lead (shown in Fig. 9). This was expected, as previous studies have shown that the extracranial trajectory of DBS systems affects their coupling with MRI electric fields [24,49,50,52,64,65,66], which in turn affects the RF-induced image artifact [49].

The overall largest distance *dz* between the actual lead tip and the artifact’s tip was 4.99 mm for trajectory 6 (T_1_MPRAGE, axial acquisition plane, right-left phase encoding direction). This distance is quite large compared to small target brain structures such as the STN, which can be around 6 × 4 × 5 mm in volume on MR images [67].

Our results challenge the paradigm in DBS lead localization that identifies the electrode tip as the center of the most caudal point within the MRI signal void, in agreement with more recent studies [33,59]. Our value of *dz* represents the difference between the actual tip of the lead and the caudal tip of the lead’s MRI artifact along the axis of the lead’s shaft. We show that this distance varies depending on the MRI sequence parameters and the patient’s extracranial trajectory. Acquisition plane and phase encoding direction had statistically significant differences in *dz*, demonstrating the importance of consistently reporting these parameters so mitigation techniques can be tailored accordingly. Therefore, for a better estimate of the depth of the DBS lead within its MRI artifact, one can reference Table 2 for the distance they should rostrally move along the lead’s shaft to identify the actual lead tip. All in all, these results will have a significant impact on the clinical interpretation of the DBS outcome and in predictions of imaging-based computational models of stimulation [20,68].

The results of this study are limited to the Siemens Aera 1.5 T scanner, the specific sequence parameters used, and unilateral DBS lead implantations on a patient’s right side. Overall, more work is required to investigate the differences in the offset of the lead’s artifact tip from the actual lead tip, including phantom experiments with the DBS lead implanted on the patient’s left side and bilaterally. Other variables that should be investigated include DBS lead manufacturer, new MRI sequences, and variation among DBS lead models, which have different contact spacing and can be directional or non-directional. Specifically, as the lead’s material and geometry of interconnecting wires can affect both the susceptibility and non-susceptibility artifact [69,70], different lead models should be studied separately.

## References

1 Deuschl G, Schade-Brittinger C, Krack P, Volkmann J, Schäfer H, Bötzel K, et al. A Randomized Trial of Deep-Brain Stimulation for Parkinson’s Disease. NEJM. 2009 Oct 8;355(9):896–908.

2 Benabid AL. Deep brain stimulation for Parkinson’s disease. Curr Opin Neurobiol. 2003 Dec 1;13(6):696–706.

3 Iorio-Morin C, Fomenko A, Kalia SK. Deep-Brain Stimulation for Essential Tremor and Other Tremor Syndromes: A Narrative Review of Current Targets and Clinical Outcomes. Brain Sci 2020, Vol 10, Page 925. 2020 Dec 1;10(12):925.

4 Zhang K, Bhatia S, Oh MY, Cohen D, Angle C, Whiting D. Long-term results of thalamic deep brain stimulation for essential tremor: Clinical article. J Neurosurg. 2010 Jun 1;112(6):1271–6.

5 Vidailhet M, Vercueil L, Houeto J-L, Krystkowiak P, Benabid A-L, Cornu P, et al. Bilateral Deep-Brain Stimulation of the Globus Pallidus in Primary Generalized Dystonia. NEJM. 2009 Oct 8;352(5):459–67.

6 Loher TJ, Capelle HH, Kaelin-Lang A, Weber S, Weigel R, Burgunder JM, et al. Deep brain stimulation for dystonia: outcome at long-term follow-up. J Neurol 2008 2556. 2008 Mar 14;255(6):881–4.

7 Perez-Malagon CD, Lopez-Gonzalez MA. Epilepsy and Deep Brain Stimulation of Anterior Thalamic Nucleus. Cureus. 2021 Sep 23;13(9).

8 Bouwens van der Vlis TAM, Schijns OEMG, Schaper FLWVJ, Hoogland G, Kubben P, Wagner L, et al. Deep brain stimulation of the anterior nucleus of the thalamus for drug-resistant epilepsy. Neurosurg Rev. 2019 Jun 1;42(2):287.

9 Baldermann JC, Melzer C, Zapf A, Kohl S, Timmermann L, Tittgemeyer M, et al. Connectivity Profile Predictive of Effective Deep Brain Stimulation in Obsessive-Compulsive Disorder. Biol Psychiatry. 2019 May 1;85(9):735–43.

10 De Koning PP, Figee M, Van Den Munckhof P, Schuurman PR, Denys D. Current status of deep brain stimulation for obsessive-compulsive disorder: A clinical review of different targets. Curr Psychiatry Rep. 2011 Apr 20;13(4):274–82.

11 Greenberg BD, Gabriels LA, Malone DA, Rezai AR, Friehs GM, Okun MS, et al. Deep brain stimulation of the ventral internal capsule/ventral striatum for obsessive-compulsive disorder: worldwide experience. Mol Psychiatry 2010 151. 2008 May 20;15(1):64–79.

12 Schlaepfer TE, Bewernick BH. Deep brain stimulation for major depression. Handb Clin Neurol. 2013 Jan 1;116:235–43.

13 Malone DA, Dougherty DD, Rezai AR, Carpenter LL, Friehs GM, Eskandar EN, et al. Deep Brain Stimulation of the Ventral Capsule/Ventral Striatum for Treatment-Resistant Depression. Biol Psychiatry. 2009 Feb 15;65(4):267–75.

14 Luo Y, Sun Y, Tian X, Zheng X, Wang X, Li W, et al. Deep Brain Stimulation for Alzheimer’s Disease: Stimulation Parameters and Potential Mechanisms of Action. Front Aging Neurosci. 2021 Mar 11;13:104.

15 Lv Q, Du A, Wei W, Li Y, Liu G, Wang XP. Deep brain stimulation: A potential treatment for dementia in Alzheimer’s disease (AD) and Parkinson’s disease dementia (PDD). Front Neurosci. 2018 May 29;12(MAY).

16 Follett KA, Weaver FM, Stern M, Hur K, Harris CL, Luo P, et al. Pallidal versus Subthalamic Deep-Brain Stimulation for Parkinson’s Disease. N Engl J Med. 2010 Jun 3;362(22):2077–91.

17 Rodriguez-Oroz MC, Obeso JA, Lang AE, Houeto JL, Pollak P, Rehncrona S, et al. Bilateral deep brain stimulation in Parkinson’s disease: a multicentre study with 4 years follow-up. Brain. 2005 Oct 1;128(10):2240–9.

18 Welter ML, Schüpbach M, Czernecki V, Karachi C, Fernandez-Vidal S, Golmard JL, et al. Optimal target localization for subthalamic stimulation in patients with Parkinson disease. Neurology. 2014 Apr 15;82(15):1352.

19 Wodarg F, Herzog J, Reese R, Falk D, Pinsker MO, Steigerwald F, et al. Stimulation site within the MRI-defined STN predicts postoperative motor outcome. Mov Disord. 2012 Jun;27(7):874–9.

20 Jiang F, Elahi B, Saxena M, Telkes I, Dimarzio M, Pilitsis JG, et al. Patient-specific modeling of the volume of tissue activated (VTA) is associated with clinical outcome of DBS in patients with an obsessive-compulsive disorder. Proc Annu Int Conf IEEE Eng Med Biol Soc EMBS. 2021;5889–92.

21 McIntyre CC, Foutz TJ. Computational modeling of deep brain stimulation. Handb Clin Neurol. 2013;116:55.

22 McIntyre CC, Mori S, Sherman DL, Thakor N V., Vitek JL. Electric field and stimulating influence generated by deep brain stimulation of the subthalamic nucleus. Clin Neurophysiol. 2004 Mar 1;115(3):589–95.

23 Volkmann J, Herzog J, Kopper F, Geuschl G. Introduction to the programming of deep brain stimulators. Mov Disord. 2002 Mar 1;17(S3):S181–7.

24 Bhusal B, Stockmann J, Guerin B, Mareyam A, Kirsch J, Wald LL, et al. Safety and image quality at 7T MRI for deep brain stimulation systems: Ex vivo study with lead-only and full-systems. PLoS One. 2021 Sep 1;16(9):e0257077.

25 Bhusal B, Keil B, Rosenow J, Kazemivalipour E, Golestanirad L. Patient’s body composition can significantly affect RF power deposition in the tissue around DBS implants: ramifications for lead management strategies and MRI field-shaping techniques. Phys Med Biol. 2021 Jan 7;66(1).

26 Golestanirad L, Keil B, Angelone LM, Bonmassar G, Mareyam A, Wald LL. Feasibility of using linearly polarized rotating birdcage transmitters and close-fitting receive arrays in MRI to reduce SAR in the vicinity of deep brain simulation implants. Magn Reson Med. 2017 Apr 1;77(4):1701–12.

27 Kazemivalipour E, Bhusal B, Vu J, Lin S, Nguyen BT, Kirsch J, et al. Vertical open-bore MRI scanners generate significantly less radiofrequency heating around implanted leads: A study of deep brain stimulation implants in 1.2T OASIS scanners versus 1.5T horizontal systems. Magn Reson Med. 2021 Sep 1;86(3):1560–72.

28 Boutet A, Chow CT, Narang K, Elias GJB, Neudorfer C, Germann J, et al. Improving Safety of MRI in Patients with Deep Brain Stimulation Devices. Radiology. 2020 Aug 1;296(2):250–62.

29 Kazemivalipour E, Keil B, Vali A, Rajan S, Elahi B, Atalar E, et al. Reconfigurable MRI technology for low-SAR imaging of deep brain stimulation at 3T: Application in bilateral leads, fully-implanted systems, and surgically modified lead trajectories. Neuroimage. 2019 Oct 1;199:18

30 Lozano CS, Ranjan M, Boutet A, Xu DS, Kucharczyk W, Fasano A, et al. Imaging alone versus microelectrode recording-guided targeting of the STN in patients with Parkinson’s disease. J Neurosurg. 2018 Jun 1;1306(6):1847–52.

31 Schenck JF. The role of magnetic susceptibility in magnetic resonance imaging: MRI magnetic compatibility of the first and second kinds. Med Phys. 1996 Jun 1;23(6):815–50.

32 Pinsker MO, Herzog J, Falk D, Volkmann J, Deuschl G, Mehdorn M. Accuracy and distortion of deep brain stimulation electrodes on postoperative MRI and CT. Zentralbl Neurochir. 2008 Aug 29;69(3):144–7.

33 Pollo C, Villemure JG, Vingerhoets F, Ghika J, Maeder P, Meuli R, et al. Magnetic resonance artifact induced by the electrode Activa 3389: an in vitro and in vivo study. Acta Neurochir 2003 1462. 2004 Dec 22;146(2):161–4.

34 Lee JY, Kim JW, Lee JY, Lim YH, Kim C, Kim DG, et al. Is MRI a reliable tool to locate the electrode after deep brain stimulation surgery? Comparison study of CT and MRI for the localization of electrodes after DBS. Acta Neurochir (Wien). 2010 Dec;152(12):2029–36.

35 Saleh C, Dooms G, Berthold C, Hertel F. Post-operative imaging in deep brain stimulation: A controversial issue. Neuroradiol J. 2016 Aug 1;29(4):244.

36 Kremer NI, Oterdoom DLM, van Laar PJ, Piña-Fuentes D, van Laar T, Drost G, et al. Accuracy of Intraoperative Computed Tomography in Deep Brain Stimulation—A Prospective Noninferiority Study. Neuromodulation Technol Neural Interface. 2019 Jun 1;22(4):472–7.

37 Bot M, Van Den Munckhof P, Bakay R, Stebbins G, Verhagen Metman L. Accuracy of Intraoperative Computed Tomography during Deep Brain Stimulation Procedures: Comparison with Postoperative Magnetic Resonance Imaging. Stereotact Funct Neurosurg. 2017 Jul 1;95(3):183.

38 Mirzadeh Z, Chapple K, Lambert M, Dhall R, Ponce FA. Validation of CT-MRI fusion for intraoperative assessment of stereotactic accuracy in DBS surgery. Mov Disord. 2014 Dec 1;29(14):1788–95.

39 Thani NB, Bala A, Swann GB, Lind CRP. Accuracy of postoperative computed tomography and magnetic resonance image fusion for assessing deep brain stimulation electrodes. Neurosurgery. 2011 Jul;69(1):207–14.

40 Girgis F, Zarabi H, Said M, Zhang L, Shahlaie K, Saez I. Comparison of Intraoperative Computed Tomography Scan with Postoperative Magnetic Resonance Imaging for Determining Deep Brain Stimulation Electrode Coordinates. World Neurosurg. 2020 Jun 1;138:e330–5.

41 Sauner D, Runge M, Poggenborg J, Maarouf M, Sturm V, Treuer H, et al. Multimodal Localization of Electrodes in Deep Brain Stimulation: Comparison of Stereotactic CT and MRI with Teleradiography. Stereotact Funct Neurosurg. 2010 Jul;88(4):253–8.

42 Engelhardt J, Guehl D, Damon-Perrière N, Branchard O, Burbaud P, Cuny E. Localization of deep brain stimulation electrode by image registration is software dependent: A comparative study between four widely used software programs. Stereotact Funct Neurosurg. 2019 Feb 1;96(6):364–9.

43 Geevarghese R, Ogorman Tuura R, Lumsden DE, Samuel M, Ashkan K. Registration Accuracy of CT/MRI Fusion for Localisation of Deep Brain Stimulation Electrode Position: An Imaging Study and Systematic Review. Stereotact Funct Neurosurg. 2016 Aug 1;94(3):159–63.

44 Hunsche S, Sauner D, Maarouf M, Poggenborg J, Lackner K, Sturm V, et al. Intraoperative X-ray detection and MRI-based quantification of brain shift effects subsequent to implantation of the first electrode in bilateral implantation of deep brain stimulation electrodes. Stereotact Funct Neurosurg. 2009;87(5):322–9.

45 Pallavaram S, Dawant BM, Remple MS, Neimat JS, Kao C, Konrad PE, et al. Effect of brain shift on the creation of functional atlases for deep brain stimulation surgery. Int J Comput Assist Radiol Surg. 2010;5(3):221.

46 Halpern CH, Danish SF, Baltuch GH, Jaggi JL. Brain Shift during Deep Brain Stimulation Surgery for Parkinson’s Disease. Stereotact Funct Neurosurg. 2008 Dec;86(1):37–43.

47 Niederer J, Patriat R, Rosenberg O, Palnitkar T, Darrow D, Park MC, et al. Factors Influencing Electrode Position and Bending of the Proximal Lead in Deep Brain Stimulation for Movement Disorders. Stereotact Funct Neurosurg. 2020 Sep 1;98(5):300–12.

48 Morishita T, Hilliard JD, Okun MS, Neal D, Nestor KA, Peace D, et al. Postoperative lead migration in deep brain stimulation surgery: Incidence, risk factors, and clinical impact. PLoS One. 2017 Sep 1;12(9):e0183711.

49 Bhusal B, Nguyen BT, Sanpitak PP, Vu J, Elahi B, Rosenow J, et al. Effect of Device Configuration and Patient’s Body Composition on the RF Heating and Nonsusceptibility Artifact of Deep Brain Stimulation Implants During MRI at 1.5T and 3T. J Magn Reson Imaging. 2021 Feb 1;53(2):599–610.

50 Vu J, Bhusal B, Rosenow J, Pilitsis J, Golestanirad L. Modifying surgical implantation of deep brain stimulation leads significantly reduces RF-induced heating during 3 T MRI. Annu Int Conf IEEE Eng Med Biol Soc IEEE Eng Med Biol Soc Annu Int Conf. 2021;2021:4978–81.

51 Sanpitak P, Bhusal B, Nguyen BT, Vu J, Chow K, Bi X, et al. On the accuracy of Tier 4 simulations to predict RF heating of wire implants during magnetic resonance imaging at 1.5 T. Annu Int Conf IEEE Eng Med Biol Soc IEEE Eng Med Biol Soc Annu Int Conf. 2021;2021:4982–5.

52 Golestanirad L, Angelone LM, Iacono MI, Katnani H, Wald LL, Bonmassar G. Local SAR near deep brain stimulation (DBS) electrodes at 64 MHz and 127 MHz: A simulation study of the effect of extracranial loops. Magn Reson Med. 2017 Oct 1;78(4):1558.

53 Fedorov A, Beichel R, Kalpathy-Cramer J, Finet J, Fillion-Robin J-C, Pujol S, et al. 3D Slicer as an Image Computing Platform for the Quantitative Imaging Network [computer program on the Internet]. Version 4.11. Magn Reson Imaging. 2012 Nov;30(9):1323–41. Available from: https://www.slicer.org/

54 McNeel, R., & others. Rhinoceros 3D [computer program on the Internet]. Version 7.0. Seattle, WA: Robert McNeel & amp, Associates; 2010. Available from: https://www.rhino3d.com/

55 Microsoft Excel [computer program on the Internet]. Version 16.59. Microsoft Corporation; 2018. Available from: https://office.microsoft.com/excel

56 R Core Team. R: A language and environment for statistical computing [computer program on the Internet]. Version 4.1.3. Vienna, Austria: R Foundation for Statistical Computing; 2021. Available from: https://www.R-project.org/.

57 RStudio Team. RStudio: Integrated Development for R [computer program on the Internet]. Version 2202.02.1. Boston, MA: RStudio, PBC; 2020. Available from: http://www.rstudio.com/.

58 Erdfelder E, FAul F, Buchner A, Lang AG. Statistical power analyses using G*Power 3.1: Tests for correlation and regression analyses. Behav Res Methods 2009 414. 2009;41(4):1149–60.

59 He C, Zhang F, Li L, Jiang C, Li L. Measurement of Lead Localization Accuracy Based on Magnetic Resonance Imaging. Front Neurosci. 2021 Dec 22;15:1613.

60 Hemm S, Coste J, Gabrillargues J, Ouchchane L, Sarry L, Caire F, et al. Contact position analysis of deep brain stimulation electrodes on post-operative CT images. Acta Neurochir (Wien). 2009 Jul 15;151(7):823–9.

61 Hyam JA, Akram H, Foltynie T, Limousin P, Hariz M, Zrinzo L. What You See Is What You Get: Lead Location Within Deep Brain Structures Is Accurately Depicted by Stereotactic Magnetic Resonance Imaging. Neurosurgery. 2015 Sep 1;11 Suppl 3:412–9.

62 Ellenbogen JR, Tuura R, Ashkan K, Ellenbogen J. Localisation of DBS Electrodes Post-Implantation, to CT or MRI? Which Is the Best Option? Stereotact Funct Neurosurg. 2018 Dec 1;96(5):347–8.

63 Li Y, Buch S, He N, Zhang C, Zhang Y, Wang T, et al. Imaging patients pre and post deep brain stimulation: Localization of the electrodes and their targets. Magn Reson Imaging. 2021 Jan 1;75:34–44.

64 Golestanirad L, Kirsch J, Bonmassar G, Downs S, Elahi B, Martin A, et al. RF-induced heating in tissue near bilateral DBS implants during MRI at 1.5 T and 3T: The role of surgical lead management. Neuroimage. 2019 Jan 1;184:566.

65 McElcheran CE, Yang B, Anderson KJT, Golestanirad L, Graham SJ. Parallel radiofrequency transmission at 3 tesla to improve safety in bilateral implanted wires in a heterogeneous model. Magn Reson Med. 2017 Dec 1;78(6):2406–15.

66 Vu J, Nguyen BT, Bhusal B, Baraboo J, Rosenow J, Bagci U, et al. Machine Learning-Based Prediction of MRI-Induced Power Absorption in the Tissue in Patients with Simplified Deep Brain Stimulation Lead Models. IEEE Trans Electromagn Compat. 2021 Oct 1;63(5):1757–66.

67 Richter EO, Hoque T, Halliday W, Lozano AM, Saint-Cyr JA. Determining the position and size of the subthalamic nucleus based on magnetic resonance imaging results in patients with advanced Parkinson disease. J Neurosurg. 2004;100(3):541–6.

68 Horn A. The impact of modern-day neuroimaging on the field of deep brain stimulation. Curr Opin Neurol. 2019 Aug 1;32(4):511–20.

69 Golestanirad L, Angelone LM, Kirsch J, Downs S, Keil B, Bonmassar G, et al. Reducing RF-induced Heating near Implanted Leads through High-Dielectric Capacitive Bleeding of Current (CBLOC). IEEE Trans Microw Theory Tech. 2019 Mar 1;67(3):1265–73.

70 Graf H, Lauer UA, Berger A, Schick F. RF artifacts caused by metallic implants or instruments which get more prominent at 3 T: an in vitro study. Magn Reson Imaging. 2005;23(3):493–9.

